# Effects of phase encoding direction on test-retest reliability of human functional connectome

**DOI:** 10.1101/2023.03.18.533301

**Authors:** Hengyi Cao, Anita D. Barber, Jose M. Rubio, Miklos Argyelan, Juan A. Gallego, Todd Lencz, Anil K. Malhotra

**Affiliations:** Institute of Behavioral Sciences, Feinstein Institutes for Medical Research, Manhasset, NY; Division of Psychiatry Research, Zucker Hillside Hospital, Glen Oaks, NY; Department of Psychiatry, Zucker School of Medicine at Hofstra/Northwell, Hempstead, NY

## Abstract

The majority of human connectome studies in the literature based on functional magnetic resonance imaging (fMRI) data use either an anterior-to-posterior (AP) or a posterior-to-anterior (PA) phase encoding direction. However, whether and how phase encoding direction would affect test-retest reliability of functional connectome is unclear. Here, in a sample of healthy subjects with two sessions of fMRI scans separated by 12 weeks (two runs per session, one with AP, the other with PA), we tested the influence of phase encoding direction on global and nodal connectivity in the constructed brain networks. All data underwent the state-of-the-art Human Connectome Project (HCP) pipeline to correct for phase-encoding-related distortions before entering analysis. We found that at the global level, the PA scans showed significantly higher intraclass correlation coefficients (ICCs) for global connectivity compared with AP scans, which was particularly prominent when using the Seitzman-300 atlas (versus the CAB-NP-718 atlas). At the nodal level, regions most strongly affected by phase encoding direction were consistently mapped to the cingulate cortex and temporal lobe, with significantly higher ICCs during PA scans compared with AP scans, regardless of atlas. Further, we demonstrated that the observed reliability differences between phase encoding directions may relate to a similar effect on the reliability of temporal signal-to-noise ratio (tSNR) in the same regions (that PA scans were associated with higher reliability of tSNR than AP scans). Averaging the connectivity outcome from the AP and PA scans could slightly, but overall have limited value to boost the ICCs. These results were largely replicated in an independent, public dataset from the HCP-Early Psychosis (HCP-EP) study with a similar design but a much shorter scan session interval. Our findings suggest that phase encoding direction has significant effects on the reliability of connectomic estimates in fMRI studies. We urge that these effects need to be carefully considered in future neuroimaging designs, especially in longitudinal studies such as those related to neurodevelopment or clinical intervention.

## Introduction

The broad application of functional magnetic resonance imaging (fMRI) in the study of human functional connectome at the macroscale has considerably promoted the understanding of the brain function and organization in healthy individuals (Power et al., 2011), neurodevelopment (Somerville et al., 2018), aging (Bookheimer et al., 2019), and among different mental disorders (Baker et al., 2019; Cao et al., 2020; Cao et al., 2021b; Dong et al., 2018; Ilioska et al., 2022; Javaheripour et al., 2021). Albeit the fruitful discoveries so far, a key consideration in the connectomic research is test-retest reliability, which quantifies the consistency of the brain connectivity readouts across multiple assessments and therefore justifies the validity and practicality of the functional connectomic measures, in particular for studies with a longitudinal design. For this purpose, a great number of studies in the literature have sought to investigate test-retest reliability of the human functional connectome (Anderson et al., 2011; Cao et al., 2019; Cao et al., 2014; Noble et al., 2019; Noble et al., 2017a; Pannunzi et al., 2017; Shah et al., 2016; Shehzad et al., 2009), and to identify factors that may help to improve the reliability of the outcome (Birn et al., 2013; Cao et al., 2019; Noble et al., 2017b; Pannunzi et al., 2017; Yoo et al., 2019). While reliability results reported in these prior studies are variable, it has generally been accepted that factors such as scan length (Anderson et al., 2011; Birn et al., 2013; Gordon et al., 2017; Noble et al., 2017b), global signal regression (GSR) (Cao et al., 2019; Noble et al., 2019; Song et al., 2012), brain atlas (Cao et al., 2019; Cao et al., 2014; Noble et al., 2019), and type of outcome measure (multivariate vs univariate)(Noble et al., 2019; Noble et al., 2017b; Pannunzi et al., 2017; Yoo et al., 2019) may play a critical role in tuning the connectome reliability. Specifically, best reliability tends to be acquired from longer scan time, without GSR, more brain nodes with finer parcellation, and the assessment of multivariate or summarized connectivity scores compared with single connectivity strength.

Phase encoding (PE) is an important technique to pinpoint the spatial location of voxel signals along the y-axis during the fMRI scans, which is achieved by applying a PE gradient to impose a specific phase angle to a transverse magnetization vector. The most frequently used PE directions (PED) in the fMRI studies are either from anterior to posterior (hereafter “AP”) or from posterior to anterior (hereafter “PA”). Different PEDs are known to have different impacts on susceptibility-induced distortion and signal loss in the brain, which may in turn influence image quality and the reliability of functional connectome constructed from these images. Specifically, previous studies have shown that AP scans are commonly associated with greater signal loss in the orbitofrontal cortex, anterior cingulate cortex (ACC), and temporal lobe (Mori et al., 2018; Wang et al., 2021; Weiskopf et al., 2006; Weiskopf et al., 2007). Less consistent findings are reported for the PA scans, which may relate to worse signals in the frontal pole, striatum, and cerebellum (Mori et al., 2018; Wang et al., 2021). Although field maps have increasingly been collected and used with the aim of correcting for PED-related distortion, the efficacy of such corrections has not been fully investigated. This raises concern as whether PED would have significant effects on test-retest reliability of the derived connectomic measures. Notably, recent studies have demonstrated significant PED-related effects on the connectivity outcome in comparison of patients with schizophrenia and healthy controls (Mori et al., 2018) as well as contrasting males and females (Wang et al., 2021), further strengthening the possibility of such concern.

When designing a neuroimaging study, investigators need to make decisions about which PED to use, despite uncertainty about how results may be affected by specific directions. As a result, many recent studies have acquired data with both directions in consecutive runs in the same session (Bookheimer et al., 2019; Demro et al., 2021; Somerville et al., 2018). Systematic differences in reliability between PEDs would impact the accuracy of the detected outcome, and therefore it might be advisable to recommend prioritizing one PED over the other when designing experiments. In this study, we examined the effects of PED on test-retest reliability of fMRI-based functional connectomic measures in healthy young adults, using two independent datasets with repeated fMRI scans. In the discovery dataset, participants received two sessions of scans with 12-week apart. The replication dataset was part of the Human Connectome Project Early Psychosis (HCP-EP) study, in which fMRI scans were acquired twice on the same day separated by approximately half an hour. During each scan session in both datasets, both AP and PA images were acquired that allows direct comparison of the reliability between the two PEDs. For each PED, we constructed connectomes using two state-of-the-art functional brain atlases (the Seitzman atlas including 300 parcels (Seitzman et al., 2020) and the CAB-NP atlas including 718 parcels (Ji et al., 2019)), both with and without GSR, and further tested if there could be any interactive effects between the PED and these different processing strategies. All data went through the Human Connectome Project (HCP) pipeline to correct for FE-related effects before entering the reliability analysis.

## Methods and Materials

### Subjects

Two independent healthy young adult datasets with repeated fMRI scans were included in the study. The discovery dataset consisted of 32 healthy participants (age 28.1±3.9 years, 13 males). The exclusion criteria included: 1) lifetime history of any psychiatric disorders as determined by Structured clinical Interview for DSM-5 (SCID), non-patient version; 2) lifetime history of neurological disorders; 3) mental retardation; and 4) MR imaging contraindications. All participants provided written informed consent for protocols approved by the Institutional Review Board of Northwell Health. The replication dataset was part of the HCP-EP study (https://www.humanconnectome.org/study/human-connectome-project-for-early-psychosis/), including 49 healthy subjects (age 24.9±4.3 years, 30 males) who provided their written consents for protocols approved by Harvard University and Indiana University.

### Study design and data acquisition

In the discovery dataset, each participant underwent two sessions of multi-paradigm fMRI scans with a time interval of 12 weeks. During each session, resting-state scans and two task-fMRI scans were performed (an event-related cognitive control task and a block-designed reward processing task). For each paradigm, two runs of data were collected with different phase encoding directions (one with AP and the other with PA) in pseudorandom order, with the sequence of the two runs counterbalanced across the sample. An overall diagram for study design is presented in Figure 1. Here, resting state included two 7-min eyes-closed runs. The cognitive control paradigm was based on the Multi-Source Interference Task (MSIT (Bush and Shin, 2006)), during which subjects were shown with three numbers (1, 2, or 3) at each trial. They were asked to select the number that is different from the other two, while ignoring the location of the number. The task lasted for a total of 8 min separated by two runs (4 min with AP and 4 min with PA). The reward processing task was similar to the one used in the Human Connectome Project (Barch et al., 2013), where subjects were asked to guess whether the number to be emerge after a question mark was above or below five during each trial. They either won or lost money based on the correctness of their responses. The task lasted for 6 min with 3-min AP scans and 3-min PA scans. Notably, all of the 32 subjects completed the resting-state scans for both sessions with 12-week apart, while only 20 and 23 subjects completed the cognitive control and reward processing tasks for both sessions, respectively. Due to differences in scan length and sample size between the three paradigms, we did not directly compare reliability outcomes between fMRI paradigms but rather treated paradigm as a random-effect variable throughout the entire study.

**Figure 1.**
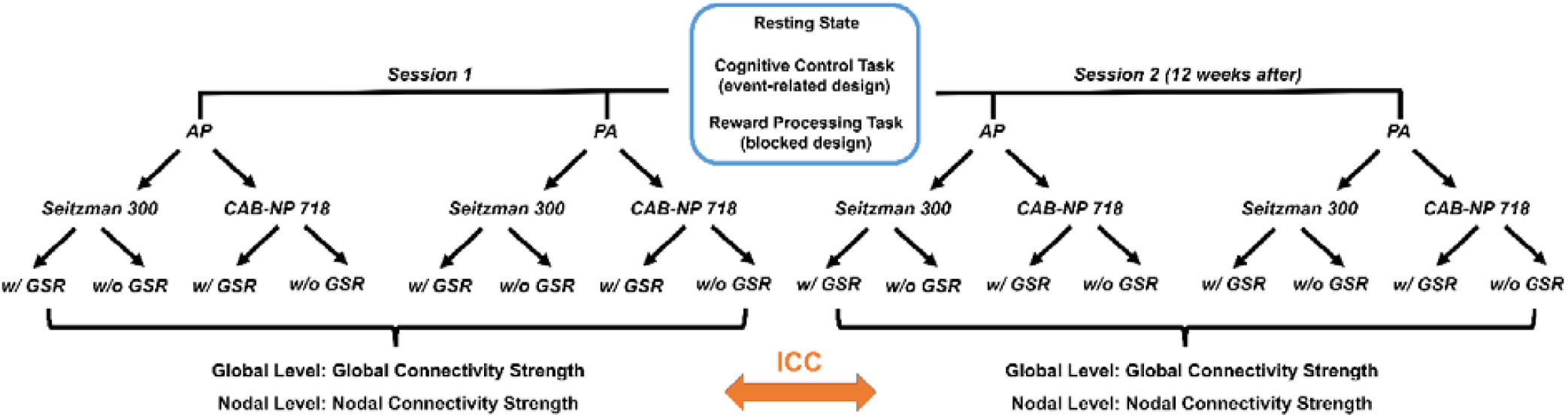
Data analysis pipeline in the discovery sample. A similar pipeline was also used for the replication sample, with data acquired from a single paradigm (resting state) and much shorter scan interval (two sessions on the same day).

In the replication dataset, two sessions of resting-state scans were conducted on the same day (separated by approximately half an hour). During each session, two runs of data were collected (one with AP, the other with PA), each lasting for 5.6 min. Notably, we used our own dataset as the discovery sample and the HCP-EP as the replication dataset since our own data were scanned 12-week apart, which resembles a typical clinical longitudinal study to investigate interventional effects. The replication sample therefore, was used to further validate whether a similar effect could be observed with a much shorter scan interval.

Imaging data from both datasets were scanned on 3T SIEMENS Prisma scanners following the HCP protocol (Van Essen et al., 2012). Specifically, the BOLD images were acquired with multi-band echo-planar imaging (EPI) sequence. Except for PED, all other parameters remained the same across all runs and sessions. The scan parameters are: 1) discovery sample: TR = 720 ms, TE = 33 ms, FA = 52 degree, slice thickness = 2 mm, 72 continuous slices, FOV = 231*231 mm, voxel size = 2.2*2.2*2 mm, multi-band factor = 8; 2) replication sample: TR = 800 ms, TE = 37 ms, FA = 52 degree, slice thickness = 2 mm, 72 continuous slices, FOV = 208*208 mm, voxel size = 2*2*2 mm, multi-band factor = 8.

### Data preprocessing

All image data were preprocessed with the HCP pipeline (Glasser et al., 2013), including a total of five major steps (PreFreeSurfer, FreeSurfer, PostFreeSurfer, fMRI Volume, fMRI Surface). Briefly, structural images were corrected for gradient- and phase encoding-related distortions, aligned at the native space, and further registered to the standard MNI space. The distortion-corrected images were submitted to the FreeSurfer *recon-all* command to segment the volume into predefined structures, reconstruct white and pial cortical surfaces, and perform FreeSurfer’s standard folding-based surface registration to their surface atlas. Similarly, functional images were first corrected for gradient and phase encoding distortions, realigned to reduce head motion, registered to the native space, and then normalized to the MNI space.

The preprocessed images were further scrutinized for head motion. In particular, we calculated the frame-wise displacements (FD) for each participant based on Power et al. (Power et al., 2012). Subjects with an average FD either > 0.5 mm or larger than the group mean plus three times the standard deviation during each paradigm were excluded for further analysis. This led to the rejection of one subject for the cognitive control task and one subject for the reward processing task in the discovery sample, as well as one subject in the replication sample.

### Construction of whole-brain connectome

Two state-of-the-art atlases covering the entire brain were used to construct functional connectome matrices (the Seitzman 300 (Seitzman et al., 2020) and the CAB-NP 718 (Ji et al., 2019)). In brief, the Seitzman-300 atlas is an updated version of the widely used Power-264 atlas (Power et al., 2011) to include the subcortex and cerebellum, and the CAB-NP-718 atlas is an extension of the Glasser’s HCP atlas (Glasser et al., 2016) to include the subcortex and cerebellum. For each node defined in these atlases, the time series were extracted and corrected for the effects of task-evoked coactivations (for task data), white matter and cerebrospinal fluid signals, 24 head motion parameters (i.e. 6 translational and rotational parameters, their first derivatives, and the squares of these 12 parameters), and FD. These noise-corrected time series were then temporally filtered (rest data: band pass 0.01-0.1 Hz; task data: high pass 0.01 Hz) and subsequently used to compute brain connectome matrices based on pairwise Pearson correlations. Notably, since whether to apply global signal regression (GSR) is still an open question (Murphy and Fox, 2017), we calculated the connectome matrices both with and without GSR in this study. The above processing pipeline was kept the same for all scan sessions and runs (Figure 1).

### Reliability assessments

The reliability of the derived connectome matrices was assessed at two levels. At the global level, we calculated the measure of global functional connectivity, which is the grand mean of the whole connectome matrices. At the nodal level, the reliability was evaluated for the connectivity strength of each single node (i.e. mean of connectivity between a given node and all other nodes in the matrices). Here, following the previous work, test-retest reliability was quantified by intraclass correlation coefficient (ICC(2,1)) based on the following formula, which gauges the absolute agreement of the measurements between two sessions.

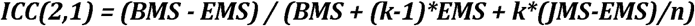

Here, *BMS* is the between-subject mean square, *EMS* is the residual mean square, *JMS* is the between-session mean square, *n* is the number of subjects, and *k* is the number of sessions. A larger positive ICC value (close to 1) indicates a better agreement of measurements between two sessions and thereby higher test-retest reliability. Based on common definition (Cao et al., 2021a; Cao et al., 2019; Cao et al., 2014), an ICC of 0.4 and above indicates fair reliability of the examined measurement, and ICC > 0.6 indicates good reliability.

### Statistics

The estimated ICC values were subsequently entered into linear mixed-effect models to determine potential effects of PED on these values. For global connectivity, PED (AP vs PA), atlas (Seitzman 300 vs CAB-NP 718), and GSR (with GSR vs without GSR) were included in the model as fixed-effect variables, with paradigm as random-effect variable. Effects of PED and interactions between PED and other fixed-effect variables (PED * atlas, PED * GSR) were estimated to decide how PED would affect the reliability of global connectivity and whether such effect would be influenced by different data processing strategies. For nodal connectivity, a similar model was employed but separated by atlas, with PED and GSR modelled as fixed-effect variables and paradigm as random-effect variable. This was applied to ICCs of each of the 300 nodes in the Seitzman atlas and each of the 718 nodes in the CAB-NP atlas.

## Results

### Reliability of global and nodal connectivity

The derived ICC values for the functional connectome at the global and nodal levels are present in Figures 2 & 3. At the global level, the ICCs of global functional connectivity ranged between 0.02 and 0.65, depending on PED, atlas, paradigm, and whether GSR was applied (Figure 2). In general, for all three examined paradigms, the lowest ICC was observed with AP scans and GSR when using the Seitzman-300 atlas, while relatively higher ICCs (>0.4) were detected with PA scans, especially when GSR was not used. An ICC of 0.4 and above was also evident for PA scans when GSR was applied, with the only exception for the cognitive control task. In addition, when using the CAB-NP-718 atlas, we also observed fair to good reliability of global connectivity with AP scans and GSR, suggesting that there may be an atlas by PED interaction on the derived reliability measures.

**Figure 2.**
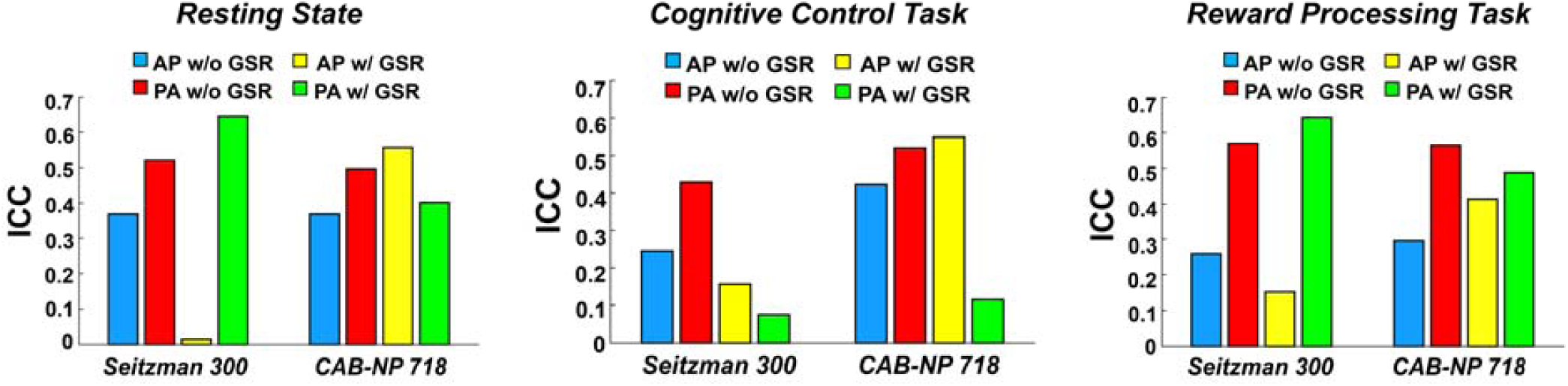
Intraclass correlation coefficients (ICCs) of global connectivity across different phase encoding directions, atlases, and global signal approaches in the discovery sample. The left, middle, and right panels represent ICCs during resting state, cognitive control, and reward processing, respectively.

**Figure 3.**
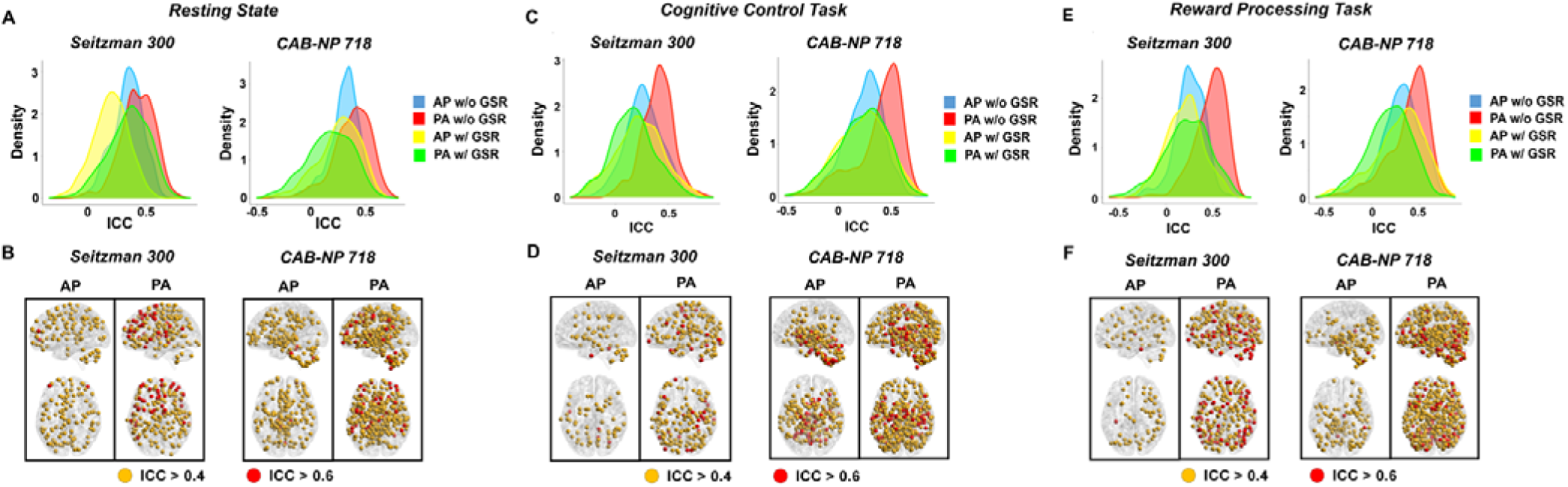
ICCs of nodal connectivity across different phase encoding directions, atlases, and global signal approaches in the discovery sample. The upper panels present the ICC distributions across all nodes in the brain, and the lower panels present the nodes with fair (ICC > 0.4, in orange) and good (ICC > 0.6, in red) test-retest reliability. The left, middle, and right panels show results during resting state, cognitive control, and reward processing, respectively. Across all paradigms and both atlases, PA scans without global signal regression had the highest reliability across the whole brain. Nodes shown on the lower panels were therefore based on ICCs without global signal regression.

In terms of functional connectivity for single nodes, relatively consistent findings were shown for both atlases and all three paradigms. Specifically, PA scans without GSR had the highest reliability, regardless of atlas and paradigm (Seitzman-300: median ICC > 0.41 for all paradigms; CAB-NP-718: median ICC > 0.40 for all paradigms), followed by AP scans without GSR (median ICC > 0.25 with Seitzman-300 and median > 0.27 with CAB-NP-718), while both AP and PA scans with GSR had relatively low reliability (median ICC > 0.17 for all paradigms with both atlases). The ICC distributions for all nodes in the constructed connectomes are present in Figure 3A, 3C, and 3E.

In Figure 3B, 3D, and 3F, we present the distribution of nodes with fair to good reliability (ICC > 0.4) in both atlases and all three paradigms. We found that during resting state, approximately 30% of total nodes in both atlases had fair reliability (ICC > 0.4) and only <1% of total nodes had good reliability (ICC > 0.6) with the AP scans. These numbers increased to >50% for nodes with fair reliability and ∼10% for nodes with good reliability with the PA scans. During the cognitive control task, fairly reliable nodes comprised of 20% and 34% of total nodes in the Seitzman-300 and CAB-NP-718 atlases respectively with the AP scans and ∼55% total nodes in both atlases with the PA scans. Similarly, proportion of nodes with good reliability increased from <7% with AP to ∼10% with PA. In high consistency, the PA scans continued to show larger proportion of nodes with fair (>60%) and good (>10%) reliability during the reward processing task, compared with 18% and <1% with the AP scans. Spatially, nodes with highest reliability during the AP scans tended to be concentrated at the posterior part of the brain, chiefly the subcortex and cerebellum; while the reliable nodes during the PA scans had a more widespread distribution across the entire brain.

### Effects of PED on reliability measures

At the global level, significant effect of PED was detected on the ICCs of global functional connectivity (*P* = 0.042), with higher reliability for PA compared with AP (Figure 4A). Moreover, there was also a significant PED by atlas interactive effect on the ICC values (*P* = 0.038), where significantly higher reliability in PA was particularly observed with the Seitzman-300 atlas (*P* = 0.028) but not with the CAB-NP-718 atlas (*P* = 0.96), suggesting that the effect of PED on reliability of global connectivity is atlas-dependent. In contrast, no significant PED by GSR interactive effect was observed.

**Figure 4.**
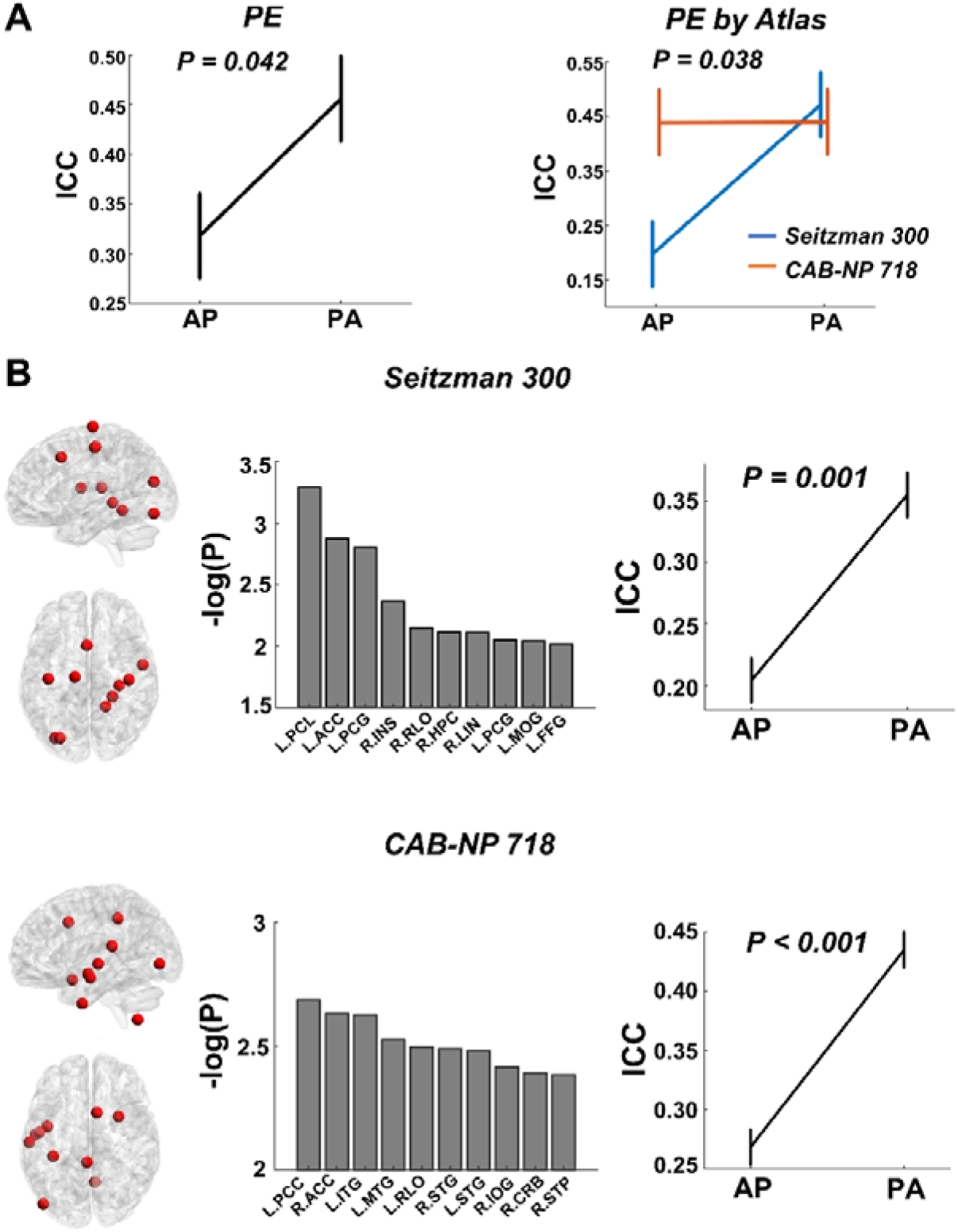
Effects of phase encoding direction (PED) on the reliability measures of global and nodal connectivity. (A) At the global level, significant PED effect and PED by atlas interactive effect were shown on the reliability of global connectivity, with reliability in PA scans significantly higher than that in AP scans, in particular using the Seitzman-300 atlas. (B) Similar effects were also observed at the nodal level, where the top ten nodes ranked by the *P* value in both atlases showed higher reliability during PA scans compared with AP scans. These nodes were largely distributed in the cingulate cortex, temporal cortex, and sensory areas with the Seitzman-300 atlas and in the cingulate cortex and temporal cortex with the CAB-NP-718 atlas. Error bars indicate standard error. Abbreviations: ACC = anterior cingulate cortex; PCL = paracentral lobule; PCG = postcentral gyrus; INS = insula; RLO = Rolandic operculum; HPC = hippocampus; LIN = lingual gyrus; MOG = middle occipital gyrus; FFG = fusiform gyrus; PCC = posterior cingulate cortex; ITG = inferior temporal gyrus; MTG = middle temporal gyrus; STG = superior temporal gyrus; IOG = inferior occipital gyrus; CRB = cerebellum; STP = superior temporal pole.

At the nodal level, ICCs of connectivity strength in 63 nodes within the Seitzman-300 atlas and in 107 nodes within the CAB-NP-718 atlas were significantly affected by PED at *P* < 0.05. While none of these effects survived false discovery rate (FDR) correction, nodes with largest effect size primarily mapped to the cingulate cortex, insula, temporal cortex, and sensory areas in the Seitzman-300 atlas and the cingulate cortex and temporal cortex in the CAB-NP atlas. The top ten nodes ranked by *P* value in each atlas are present in Figure 4B. Notably, for all of the ten top nodes in both atlases, their ICCs were significantly higher in PA compared with AP (Seitzman-300: *P* = 0.001; CAB-NP-718: *P* < 0.001), suggesting that PA scans may generally boost the reliability of functional connectivity at the nodal level independently of atlas.

### Are PED effects on reliability of connectivity related to PED effects on reliability of signal-to-noise ratio and head motion?

Based on these findings, we further asked the questions as whether the observed PED effects on reliability of functional connectivity may relate to 1) a similar PED effect on reliability of temporal signal-to-noise ratio (tSNR) in the same regions; and 2) a similar PED effect on reliability of head motion. To this end, we calculated node-specific tSNRs and FDs for the most affected nodes as shown in Figure 4. The node-specific FDs were generated based on previous publications (Satterthwaite et al., 2013; Yan et al., 2013) using the DPARSF toolbox (Chao-Gan and Yu-Feng, 2010). Briefly, the position of each voxel in the brain at each time point was estimated by applying the motion transformation matrix to the original position of each voxel derived from the reference image during realignment (here the single-band images during the scan), resulting in a series of maps quantifying the distance change of each voxel relative to its preceding time point. The node-specific FDs were further calculated by averaging all voxels within a given node. Similarly, in terms of prior definition (Chen et al., 2020; Demetriou et al., 2018), node-specific tSNRs were evaluated as the mean of the time series divided by its standard deviation in each node.

Using the same measure of ICC(2,1), we subsequently estimated the reliability of FD and tSNR for each of the top ten nodes with the strongest PED effect. The derived ICC values were then entered into a similar linear mixed model as dependent variable, with PED and atlas as fixed variables and paradigm as random variable. Here, we observed highly significant effect of PED on the reliability of tSNR across the same regions (*P* < 0.001, Figure 5A), with much higher ICCs during the PA scans compared with the AP scans. By contrast, the reliability of FD in these regions did not significantly differ between the AP and PA scans (*P* = 0.47, Figure 5B), suggesting that the detected PED effect on reliability of nodal connectivity may to certain degree relate to a similar effect on reliability of signal intensity in the same regions.

**Figure 5.**
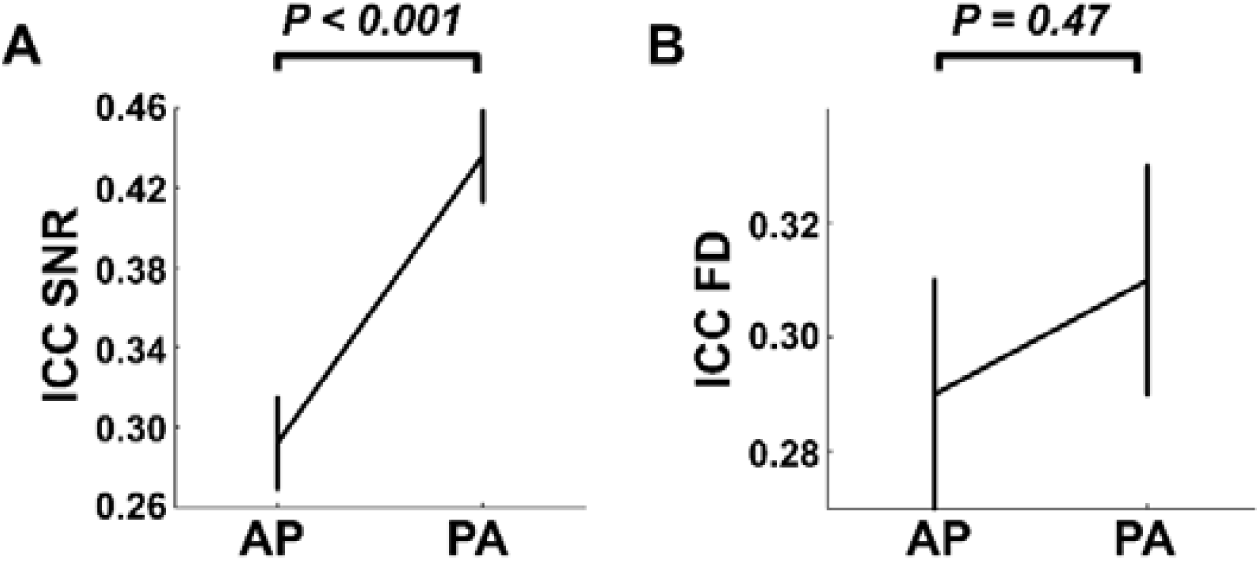
Effects of phase encoding direction on the reliabilities of signal-to-noise ratio (SNR, Panel A) and head motion (Panel B). For the same regions shown in Figure 4B, their SNR reliability was significantly higher during PA scans compared with AP scans. No significant effect was shown for the reliability of head motion. Error bars indicate standard error.

### Does averaging AP and PA scans boost the reliability?

One commonly used approach to combining data acquired from different PEDs, as recommended by the HCP, is to average the derived AP and PA functional connectivity matrices (Smith et al., 2013). This raises the question as whether test-retest reliability could be boosted by using these averaged connectivity measures. To answer this question, we further calculated the ICC(2,1) values for the averaged connectome, at the global level as well as at the nodal level particularly for nodes with the largest PED effects. As presented in Figure 6, at the global level, no significant reliability differences were shown between the PA and the averaged connectomes, regardless of atlas (*P* > 0.75). Similar findings also applied to the nodal level (*P* > 0.06), although there appeared to be a trendy effect towards higher reliability in the averaged connectome, such that ICC_Average_ > ICC_PA_ > ICC_AP_. These results suggest that while averaging AP and PA scans may slightly boost the reliability, the improvement is very limited and non-significant.

**Figure 6.**
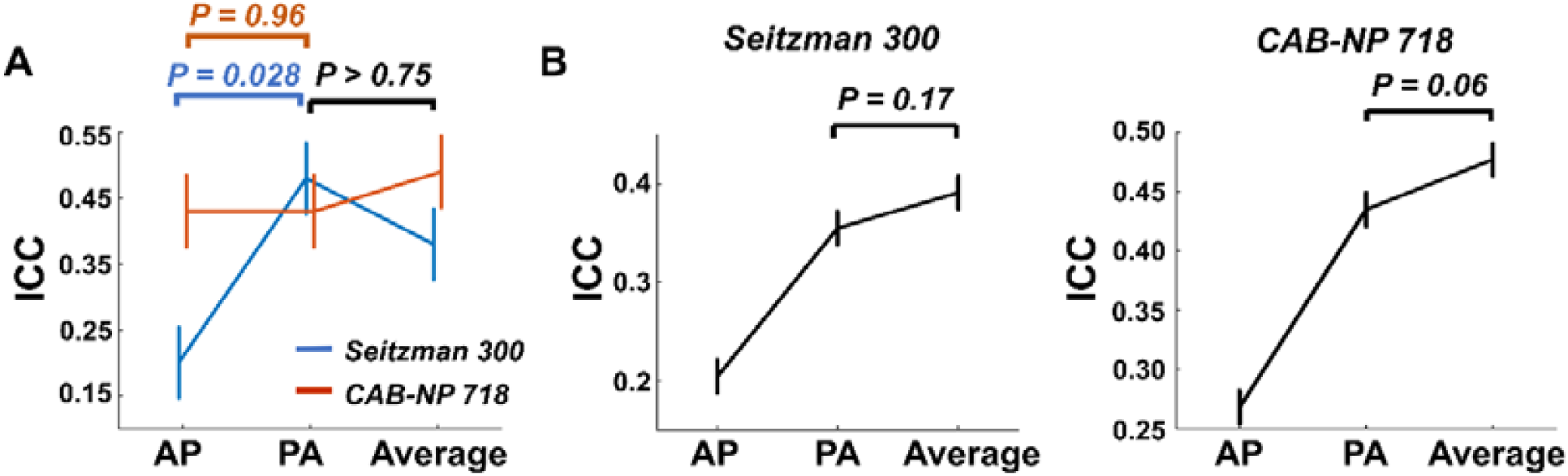
Reliability of global (Panel A) and nodal (Panel B) connectivity for data averaged by AP and PA scans. No significant reliability differences were shown between the PA and the averaged data, despite a trend-level effect towards slightly better reliability of nodal connectivity in the averaged data. Error bars indicate standard error.

### Replication of reliability findings in an independent dataset

When analyzing data from the HCP-EP project, we found overall very similar effects at both global and nodal levels. As presented in Figure 7A, at the global level, significant PED by atlas interactive effect was shown on the reliability of global connectivity, with significantly higher ICCs during PA scans with the Seitzman-300 atlas (*P* = 0.02). No significant difference was observed in ICCs between the PA scans and the average of AP and PA scans (*P* > 0.14), despite a slightly upward trend in the reliability of averaged scans. At the nodal level (Figure 7B & 7C), highly consonant with findings in the discovery sample, the top ten nodes most affected by PED showed significantly higher ICCs in PA scans compared with AP scans (Seitzman-300: *P* = 0.026; CAB-NP-718: *P* = 0.002), while non-significant but trendy effects were present between the reliability of PA scans and that of the averaged scans (*P* > 0.19). Moreover, while the exact location of the top ten nodes differed between the two samples, these nodes were nevertheless similarly distributed in the cingulate cortex, temporal cortex, insula, and sensory areas in the Seitzman-300 atlas and in the cingulate cortex, temporal cortex, cerebellum, and prefrontal cortex in the CAB-NP-718 atlas.

**Figure 7.**
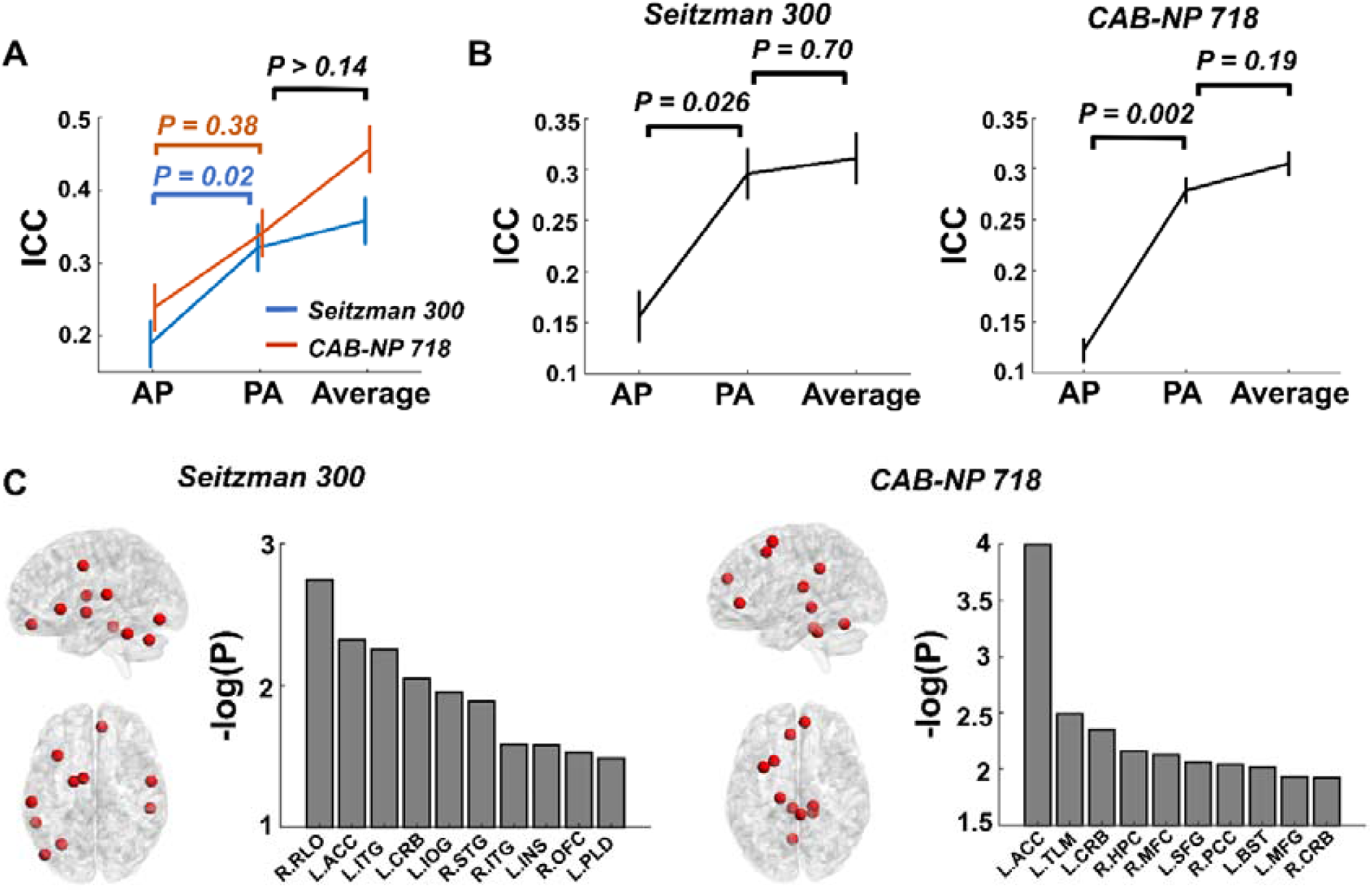
Replication of findings in the HCP-EP dataset. (A) Similar PED and PED by atlas effects were shown for reliability of global connectivity. (B) Similar PED effects on reliability of nodal connectivity were observed. (C) The most affected nodes in the replication sample were chiefly mapped to the cingulate cortex, temporal cortex, prefrontal cortex, and cerebellum, which is similar to the discovery sample. Error bars indicate standard error. Abbreviations: ACC = anterior cingulate cortex; INS = insula; RLO = Rolandic operculum; HPC = hippocampus; PCC = posterior cingulate cortex; ITG = inferior temporal gyrus; STG = superior temporal gyrus; IOG = inferior occipital gyrus; CRB = cerebellum; OFC = orbitofrontal cortex; PLD = pallidum; TLM = thalamus; MFC = medial frontal cortex; SFG = superior frontal gyrus; MFG = middle frontal gyrus; BST = brain stem.

## Discussion

This study for the first time systematically examined the effects of PED on test-retest reliability of human functional connectome. In two independent datasets with repeated scans using both AP and PA directions, we showed that PA scans were associated with significantly higher reliability of global connectivity compared with the AP scans, in particular using the Seitzman-300 atlas with relatively fewer brain nodes (versus the CAB-NP-718 atlas). At the nodal level, while reliability differed node by node, regions most strongly affected by PED were consistently mapped to the cingulate cortex and temporal cortex in both atlases, with almost uniformly increased reliability during PA scans in these regions. Further, we found that differences in test-retest reliability between different PEDs may relate to a similar PED effect on the reliability of image tSNRs, and averaging AP and PA scans may slightly, but overall have limited value to increase the reliability of functional connectivity. These findings suggest that PED has significant effects on the reliability of human functional connectome and should be taken into consideration during the design of imaging protocol, especially for longitudinal studies.

For both global and nodal connectivity, the reliability varied between different PEDs, atlases, and global signal approaches. While the effects of PED are a new finding in the present study, the other factors have been well-known to play an important role in modulating the reliability outcome in the functional connectome data. In particular, data without GSR have repeatedly been found to be more reliable than data with GSR (Cao et al., 2019; Noble et al., 2019; Song et al., 2012; Tozzi et al., 2020), and data constructed from a more fine-grained atlas with more functional parcels are frequently reported to be more reliable than those constructed from an atlas with smaller number of parcels (Cao et al., 2021a; Cao et al., 2019; Cao et al., 2014; Noble et al., 2019; Tozzi et al., 2020). Our present findings are highly consistent with these prior observations: when using the CAB-NP-718 atlas without GSR, fair to good reliability (ICC > 0.4) was detected at the global level regardless of PED and paradigm; at the nodal level, averagely ∼40% of total nodes showed fair to good reliability, across both PEDs and all paradigms. While the effects of atlas and GSR are not the research focus in the present study, we nevertheless tested the potential interactive effect between PED and these factors on the functional connectivity measures. The significant PED by atlas interaction observed in the present work implies that a more fine-grained atlas may at least partly compensate for the PED effects on the functional connectome. Therefore, these findings may together suggest that irrespective of the PED choice, the use of atlases with more functional details and no conduction of GSR are generally preferred, if test-retest reliability is the major consideration of the study.

During the AP scans, nodes with good reliability (ICC > 0.6) were largely distributed at the posterior part of the brain, chiefly the subcortex and cerebellum; while the distribution of the most reliable nodes was more widespread across the whole brain during the PA scans. The more constrained distribution of the reliable nodes during AP scans may relate to artifacts and signal distortions induced by the sinuses and eyes, which render relatively lower reliability in image quality for the anterior part of the brain compared with the posterior part. This explanation is supported by the subsequent finding that AP and PA images were associated with different reliability in tSNR, in particular regions whose reliability of connectivity was most affected by PED. One possibility that the subcortical regions are more resilient to this effect is that these regions have extensive connections with the entire cerebral cortex (both anterior and posterior part), and therefore the effects of less reliable connections at the anterior part have been cancelled out by the effects of more reliable connections at the posterior part, rendering the reliability of overall subcortical connectivity relatively stable. Statistically, regions that were consistently influenced by PED in both atlases were the ACC and temporal cortex. This is remarkably similar to results in previous studies that both functional connectivity strength and amplitude of low-frequency fluctuation (ALFF) in these regions are strongly affected by PED (Mori et al., 2018; Wang et al., 2021). Notably, these regions are not only along the PE axis during imaging acquisition but also close to air/tissue interface, which possibly suffer magnetic susceptibility induced image distortion and propagated flow or motion artifacts (De Panfilis and Schwarzbauer, 2005; Weiskopf et al., 2006). Tentatively, such effects may be more variable and less stable with the AP scans, leading to lower reliability of functional connectivity in these regions. Other regions such as the orbitofrontal cortex, posterior cingulate cortex, and sensory areas may bear a similar problem, but the effects may be more prominent when processed with certain atlases.

In this study, we tested two possibilities that may underlie the observed reliability differences between PEDs. Parallel to the interpretations above, we found that image tSNR was also more reliable during PA scans compared with the AP scans, in particular in regions with the largest PED effects on reliability of functional connectivity. This finding suggests that even after careful correction of PE effects during data processing (such as the standard pipeline implemented in the HCP), the PE effects are still not able to be fully removed and could impact the image signal intensity and stability. By contrast, no significant differences were observed between AP and PA scans in terms of reliability of head motion. As head motion is a critical consideration in the calculation of functional connectivity (Power et al., 2012; Satterthwaite et al., 2013; Yan et al., 2013) and may affect the reliability of connectivity measure (Cao et al., 2019; Noble et al., 2019; Yan et al., 2013), this result greatly mitigates the possibility that the PED effects on reliability of human functional connectome are due to differences in reliability of head motion. This is not surprising since head motion has been proposed to reflect an individualized trait (Couvy-Duchesne et al., 2014; Engelhardt et al., 2017; Zeng et al., 2014) and relate to psychopathology (Couvy-Duchesne et al., 2016; Kong et al., 2014). Therefore, head motion assessments should theoretically be associated with inter-subject variation rather than variation in scan protocol.

Our results also demonstrated that by averaging data collected from both PEDs, the reliability may slightly but non-significantly increase, suggesting that the use of averaged data may to certain degree be beneficial, yet the improvement is very limited. One possibility is that the averaged data are blended with information derived from both relatively less reliable AP scans and relatively more reliable PA scans, diluting the overall reliability in the averaged connectivity measures. However, although the effect is not statistically significant, the trends towards higher reliability in the averaged data still imply that the approach to averaging AP and PA scans is generally preferred if multiple runs of data are available (even with different PEDs), compared with using data from a single run. It remains to be determined, however, whether acquisition of multiple runs of only PA scans would further boost the reliability (versus multiple runs with different PEDs).

One major limitation of this study, as we clearly acknowledge here, is that the scan length is relatively short for each run (7 min for the discovery sample and 5.6 min for the replication sample). As prior work has shown that at least 20-30 min scan time is required to achieve a stable result (Gordon et al., 2017; Laumann et al., 2015), we cannot exclude the possibility that the findings reported here are influenced by scan length. It is important to further investigate whether longer scans would compensate for the observed effects of PED in future studies. The second limitation relates to the sample size, which is relatively small in both samples. Although similar PED effects were observed across both samples, future studies with larger test-retest datasets are still warranted to verify these findings. Thirdly, our results were purely based on the PE correction approach implemented in the HCP pipeline using field maps, which may not be generalizable to other PE correction approaches (Gu and Eklund, 2019). We employed the HCP pipeline since this is considered as a state-of-the-art approach for imaging preprocessing and has widely been used in the neuroimaging community. However, our results may not necessarily imply a lack of efficacy for the employed approach in PE correction but may reflect an intrinsic problem for the MRI scans. Fourthly, all data in both samples were collected from scanners with the same make and model (SIEMENS Prisma) using multiband EPI scans, and therefore whether the detected effects are specific to scanner and/or scan sequence needs to be examined. Finally, as both scan length and sample size differed between fMRI paradigms, we treated paradigm as a random-effect variable in this study. However, since paradigm itself may affect reliability of functional connectivity (Cao et al., 2019; Cao et al., 2014; Shah et al., 2016), direct comparison of PE effects between different paradigms should also be performed in the future.

In sum, this study for the first time reveals the effects of PED on test-retest reliability of human connectomic measures derived from fMRI data, at both global and nodal levels. Overall, the PA scans appear to show superiority in reliability of both global and nodal connectivity, suggesting that PA scans may be preferred if the research is not focused on a specific region of interest. We urge that the PED effects need to be carefully considered in future neuroimaging designs, especially in longitudinal studies such as those related to neurodevelopment or clinical intervention.

